# An Open Competition for Biomarkers of Aging

**DOI:** 10.1101/2024.10.29.620782

**Authors:** Kejun Ying, Seth Paulson, Julian Reinhard, Lucas Paulo de Lima Camillo, Jakob Träuble, Stefan Jokiel, Biomarkers of Aging Consortium, Dane Gobel, Chiara Herzog, Jesse R. Poganik, Mahdi Moqri, Vadim N. Gladyshev

**Affiliations:** Division of Genetics, Department of Medicine, Brigham and Women’s Hospital and Harvard Medical School, Boston, MA, USA; T. H. Chan School of Public Health, Harvard University, Boston, MA, USA; Methuselah Foundation, Washington, DC, USA; Global Bioinformatics, Evotec SE, Hamburg, Germany; School of Clinical Medicine, University of Cambridge, Cambridge, UK; Department of Chemical Engineering and Biotechnology, University of Cambridge, Cambridge, UK; Faculty of Physics, Ludwig Maximilian University of Munich, Munich, Germany; Institute for Biomedical Ageing Research, Universität Innsbruck and European Translational Oncology Prevention and Screening Centre, Universität Innsbruck, Innsbruck, Austria; Department of Genetics, Stanford University, Stanford, CA, USA

## Abstract

Open scientific competitions have successfully driven biomedical advances but remain underutilized in aging research, where biological complexity and heterogeneity require methodological innovations. Here, we present the results from Phase I of the Biomarkers of Aging Challenge, an open competition designed to drive innovation in aging biomarker development and validation. The challenge leverages a unique DNA methylation dataset and aging outcomes from 500 individuals, aged 18 to 99. Participants are asked to develop novel models to predict chronological age, mortality, and multi-morbidity. Results from the chronological age prediction phase show important advances in biomarker accuracy and innovation compared to existing models. The winning models feature improved predictive power and employ advanced machine learning techniques, innovative data preprocessing, and the integration of biological knowledge. These approaches have led to the identification of novel age-associated methylation sites and patterns. This challenge establishes a paradigm for collaborative aging biomarker development, potentially accelerating the discovery of clinically relevant predictors of aging-related outcomes. This supports personalized medicine, clinical trial design, and the broader field of geroscience, paving the way for more targeted and effective longevity interventions.

## Introduction

The global demographic shift towards an aging population poses unprecedented challenges to healthcare systems and societal structures^1,2^. This transition, marked by a surge in age-related pathologies, underscores the critical need for precise quantification of biological aging^3^. Chronological age fails to capture the heterogeneity in aging trajectories among individuals, requiring robust biomarkers of aging—quantifiable indicators that accurately reflect biological age and predict age-associated decline^4^.

Recent breakthroughs have revolutionized our understanding of aging biomarkers, particularly in the realm of epigenetics. The field has seen a proliferation of epigenetic clocks, including the seminal contributions of Horvath^5^, Hannum^6^, PhenoAge^7^, GrimAge^8^, and DunedinPACE^9^. More recent developments include causality-enriched clocks^10^, genomic-region-based clocks (including PRC2 clock^11^, RetroAge^12^), and deep learning-based clocks^13–15^. These advances, alongside the establishment of the Biomarkers of Aging Consortium, have laid a foundation for the identification and validation of targeted interventions to extend healthy lifespan by fostering collaboration and standardization efforts^4,16,17^. However, despite these strides, significant challenges remain, such as limited reproducibility of biomarker results across populations, insufficient longitudinal validation, and unclear mechanistic links to fundamental aging processes.

Open science initiatives have catalyzed breakthroughs in complex biomedical fields. For instance, the Critical Assessment of protein Structure Prediction (CASP) competitions dramatically advanced protein folding predictions^18^ and led to the development of Rosetta^19^ and Alphafold^20^, which have revolutionized the field of structural biology. Similarly, the DREAM Challenges advanced our understanding of gene regulatory networks and drug sensitivity^21^, and the Alzheimer’s Disease Neuroimaging Initiative (ADNI) in neurology accelerated biomarker discovery through open data and competitions^22^. In these initiatives, researchers compete to develop the best solutions while simultaneously sharing data, methods, and insights, fostering a collaborative environment that accelerates scientific progress. This unique blend of competition and cooperation has proven highly effective in tackling complex biomedical challenges. These successes underscore the immense potential of open competitions to drive rapid advancements in aging research.

Here, we present Phase I of the Biomarkers of Aging Challenge, a global open science initiative that aims to advance the development and validation of aging biomarkers.. This challenge leverages crowdsourcing and standardized datasets to overcome key barriers in the field. (1). We introduce a unique, curated dataset of DNA methylation profiles and comprehensive health outcomes from 500 individuals (ages 18-99), facilitating direct comparisons of biomarker models across diverse genetic backgrounds and environmental exposures^23^. (2). The Biomarkers of Aging Challenge utilizes Biolearn (https://bio-learn.github.io), an open-source computational platform that standardizes the implementation and evaluation of aging biomarkers^17^. Biolearn bridges the methodological gap between computational biology and data science, enabling systematic benchmarking of novel algorithms. (3). Our three-phase competition structure—encompassing chronological age prediction, mortality risk assessment, and multi-morbidity forecasting— incentivizes the development of biomarkers with direct clinical relevance, aligning with recent frameworks for evaluating longevity interventions^4^.

Preliminary findings from the chronological age prediction phase have yielded unexpected insights into the epigenetic landscapes of aging. Top-performing models have identified novel DNA methylation signatures that demonstrate superior accuracy and generalizability compared to existing biomarkers. The Biomarkers of Aging Challenge exemplifies the transformative potential of open competitions in accelerating scientific discovery. By fostering interdisciplinary collaboration and providing a unified framework for biomarker evaluation, we have created an ecosystem that rapidly translates diverse expertise into tangible progress. This approach not only addresses the heterogeneity in biomarker formulations but also tackles the inconsistencies in dataset structures that have hindered progress in the field.

## Results

### Overview of the Challenge series

The Biomarkers of Aging Challenge was designed as a multi-phase open science initiative to address key limitations in aging biomarker development and validation. Launched in March 2024, the competition attracted 152 teams from 28 countries, representing a diverse array of expertise in computational biology, data science, and geroscience (Figure 1).

**Figure 1.**
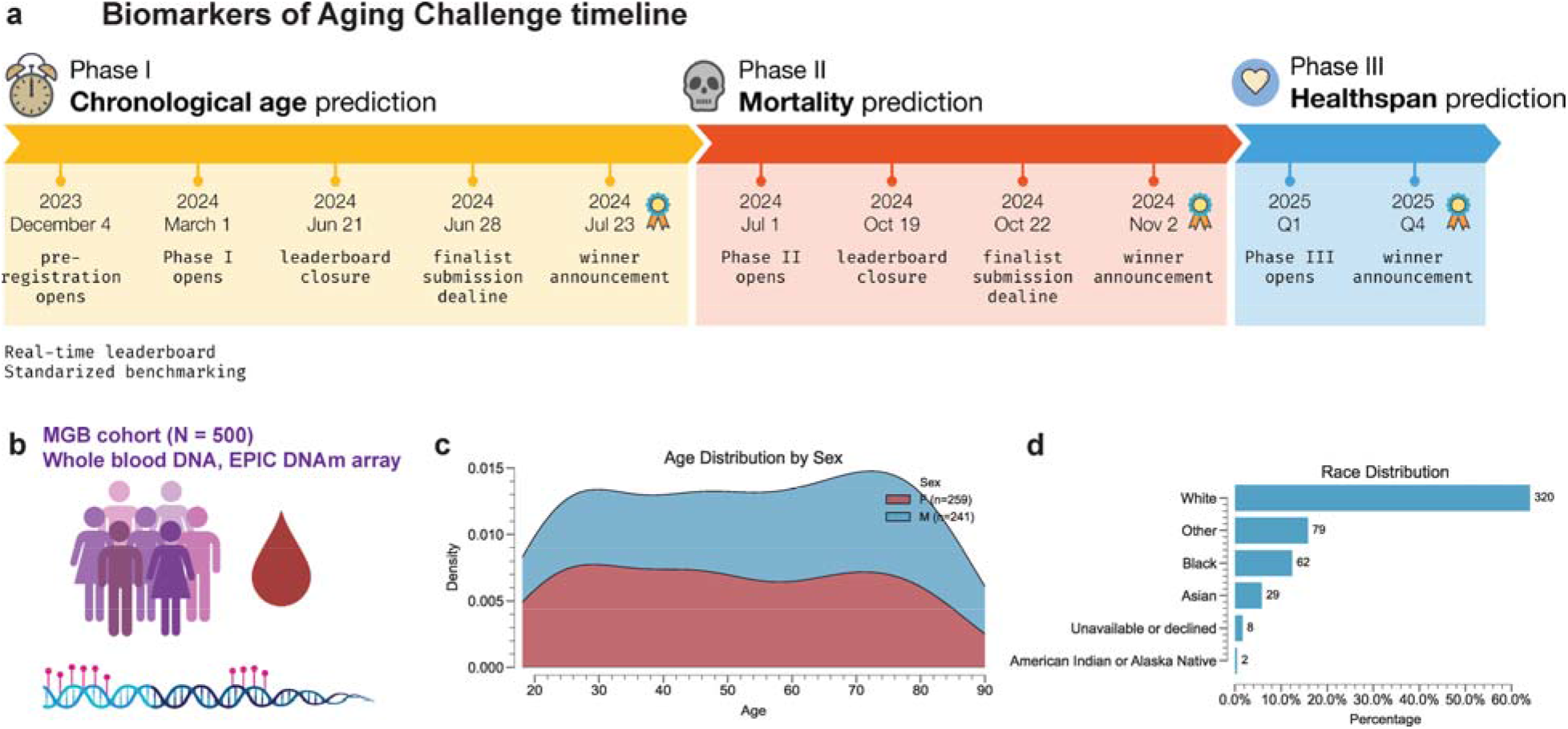
Overview of the Biomarkers of Aging Challenge. **a**. Timeline of the challenge showing the three phases: Chronological Age Prediction (March-July 2024), Mortality Prediction (July-November 2024), and Multi-morbidity Prediction (planned for 2025). Key dates for each phase, including announcements, deadlines, and result releases, are indicated. **b**. Illustration of the MGB cohort (n = 500) used in the challenge, showing blood samples, DNA extraction, and methylation array analysis. **c**. Age distribution of the challenge cohort, with density plots colored for males and females. **d**. Bar chart showing the race distribution of the challenge cohort, including White, Black, Asian, and other categories.

The challenge is structured in three sequential phases (Figure 1a):

1. Chronological Age Prediction (March - July 2024): Participants developed models to predict chronological age from DNA methylation data. This phase served as a benchmark for feature selection and model performance.
2. Mortality Prediction (July - November 2024): Teams refined their models to predict all-cause mortality risk, a key outcome in aging research.
3. Multi-morbidity Prediction (Scheduled for 2025): This phase will focus on predicting the onset of multiple age-related conditions, which will be critical for the optimization and extension of healthspan.

Central to the challenge is a unique, high-quality reference dataset. We generated methylation profiles for 500 individuals from the Mass General Brigham Biobank using the Illumina MethylationEPIC v2.0 platform, which captures methylation status at over 930,000 CpG sites^23^. This dataset covers a wide range of ages, is balanced between males and females, and represents the racial and ethnic diversity of the Boston area (Figure 1b-d). Participants were provided with the complete epigenetic dataset but were blinded to all outcome data. Model performance was evaluated using a split-sample approach: 50% of the data was used for ongoing assessment via a public leaderboard, while the remaining 50% was withheld as a final validation set. This methodology facilitated iterative model refinement while mitigating the risk of overfitting.

To standardize biomarker implementation and evaluation, we utilized our recently developed open-source Python-based Biolearn platform. Biolearn facilitated the integration of 17 additional public datasets from the Gene Expression Omnibus, encompassing 12,463 samples across diverse tissue types and age ranges (Figure 1b). This comprehensive framework enabled systematic benchmarking of both existing and novel biomarker algorithms.

Submissions were evaluated using a standardized pipeline on a held-out test set. For the chronological age prediction phase, performance was assessed using mean absolute error (MAE) and feature number. Participants were able to submit once daily, and the leaderboard was updated in real time, fostering competition and rapid iteration of models.

### Phase I: Chronological age prediction

The inaugural phase of the Biomarkers of Aging Challenge, focusing on chronological age prediction, attracted 37 teams and garnered 551 submissions. The top-performing models consistently surpassed existing published biomarkers of aging (BoAs) in chronological age prediction accuracy on our 500-sample dataset (Figure 2a). The phase I first-ranked entry (DarthVenter) achieved a mean absolute error (MAE) of 2.45 years on the final dataset, with a leaderboard score of 2.11 years. The second-ranked entry (Lucascamillo) and third-ranked entry (ZetaPartition) followed closely with an MAE of 2.55 years and 2.46 years, respectively.

**Figure 2.**
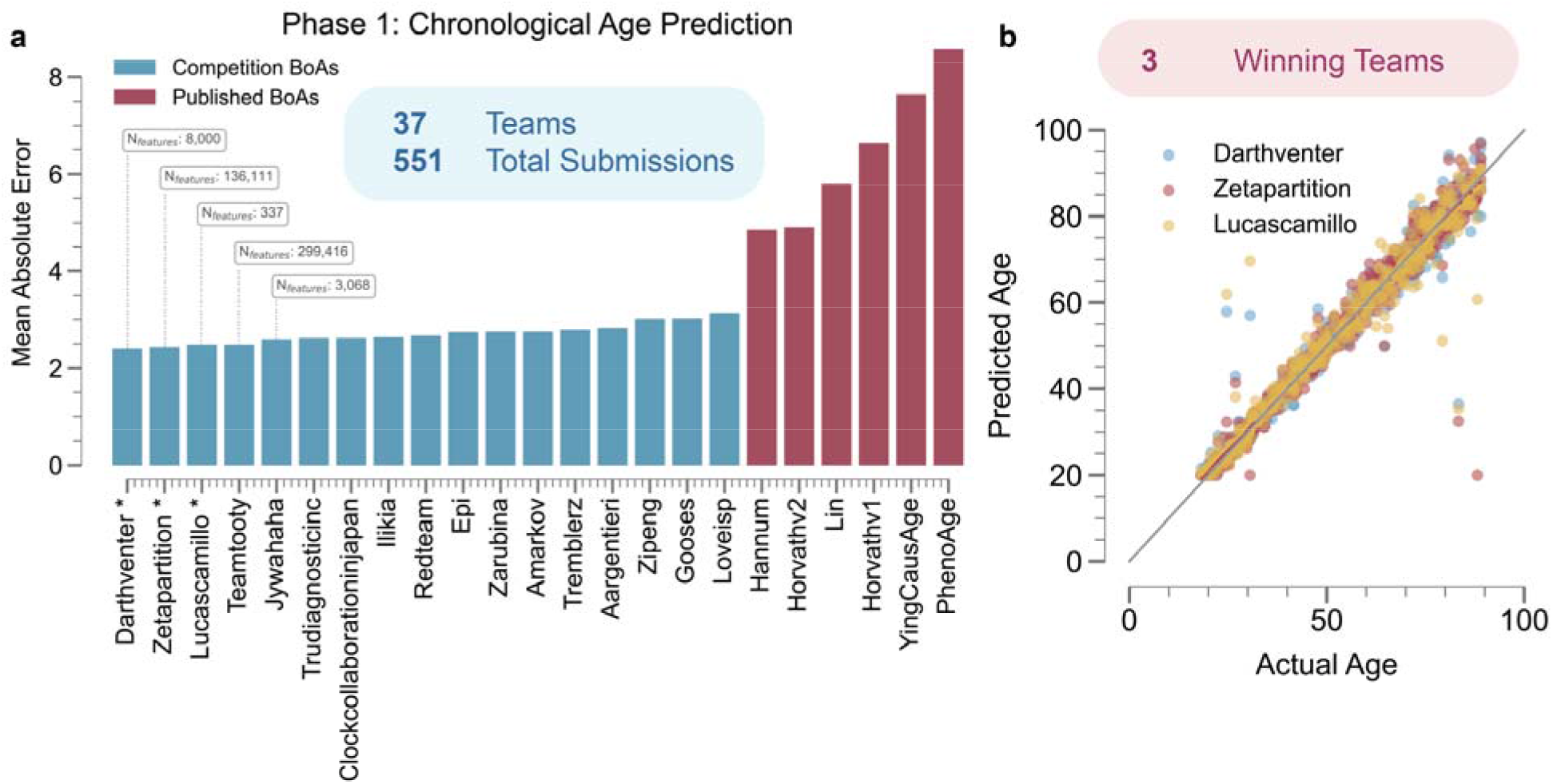
Phase 1 of the Biomarkers of Aging Challenge. **a**. Bar plot showing the mean absolute error (MAE) for different models, including competition submissions (blue) and published biomarkers (red). The x-axis lists the models, and the y-axis shows the MAE in years. The plot indicates that 37 teams submitted a total of 551 entries. **b**. Scatter plot comparing predicted vs. actual age for the top 3 winning teams. Each point represents an individual, with different colors for each team’s model. The diagonal line represents perfect prediction.

Notably, these top-performing teams employed diverse approaches in terms of feature selection. The first-ranked entry utilized 8,000 features, and the third-ranked entry leveraged 136,111 features, in contrast to the second-ranked entry’s more parsimonious approach with 337 features. This variation in feature set sizes demonstrates that high performance can be achieved through different strategies in feature selection and model complexity.

All finalist competition entries consistently achieved MAEs below 3 years, representing a significant improvement over existing biomarkers. In comparison, the best-performing published biomarker in the current Biolearn collection (Horvath) exhibited an MAE of approximately 4.8 years, with other established biomarkers such as Hannum, PhenoAge, and GrimAge demonstrating progressively higher MAEs ranging from about 5 to 8.5 years. This substantial improvement in accuracy underscores the potential of new approaches developed during the challenge to advance the field of epigenetic age prediction and offers new avenues for investigating the biological underpinnings of aging.

The winning models also employed diverse approaches (Figure 2b), showcasing the potential for innovation in epigenetic age prediction. The first-ranked entry, Skip-Improved Training Hive (SITH) network, utilized an ensemble of feed-forward neural networks with a linear skip layer, trained on 8,000 CpG sites from 19 datasets. The second-ranked entry introduced a transformer-based foundation model pre-trained on over 100,000 samples and fine-tuned for age prediction using a subset of 337 CpG sites. The third-ranked entry is a deep learning model based on a ResNet architecture, leveraging 136,111 CpG sites from 33 publicly available datasets.

Overall, these results represent a significant improvement in chronological age prediction using epigenetic data. The competition has produced models that substantially outperform existing published biomarkers, offering more accurate tools for assessing biological age. Moreover, the consistent performance across age ranges suggests robust models that could be applicable in various research and potentially clinical contexts. The outcomes of this initial phase set a new benchmark for epigenetic age prediction and provide a strong foundation for subsequent phases of the Biomarkers of Aging Challenge, which will focus on more complex outcomes such as mortality and multi-morbidity prediction.

## Discussion

The Biomarkers of Aging Consortium broadly aims to catalyze progress in the complex field of aging. We hypothesized that leveraging the power of an open science initiative, the Biomarkers of Aging Challenge Series, would yield significant advances in aging biomarker development. The results of the first phase of our challenge provide powerful proof of concept for this idea. The winning models not only outperformed existing epigenetic clocks but also introduced novel approaches that may have broader implications for aging research and personalized medicine.

The top-performing models employed diverse strategies to improve age prediction accuracy. The first-ranked entry’s SITH network utilized an ensemble method with a skip-layer architecture, which showed effectiveness in capturing age-related signals while addressing batch effects. The second-ranked entry’s CpGPT model applied a transformer-based approach, demonstrating the applicability of transfer learning in epigenetics. The models achieved MAEs ranging from 2.45 to 2.55 years, representing an improvement over existing biomarkers. The third-ranked entry’s approach incorporated a larger number of CpG sites (136,111) compared to traditional epigenetic clocks, potentially leveraging previously unexplored genomic regions. This increased precision may enhance the detection of subtle age-related changes and improve the evaluation of interventions targeting the aging process. However, the complexity of these models presents challenges in interpreting their biological significance and understanding the mechanisms underlying their predictions–an ongoing general issue in this field. Further research is needed to elucidate the biological relevance of these computational advancements and their potential applications in aging research and clinical settings.

One key challenge moving forward will be to bridge the gap between predictive accuracy and biological insight. While these models excel at chronological age prediction, their complexity may obscure the underlying biological processes they capture. Future research should focus on developing interpretable models or methods to extract biological meaning from these high-performing but opaque predictors.

The success of this challenge also highlights the value of standardized datasets and evaluation frameworks in advancing the field. The novel dataset generated for this competition, covering a wide age range and balanced for sex, provides a valuable resource for future research. The opensource Biolearn platform offers a standardized implementation of existing biomarkers and evaluation tools, which could facilitate more robust comparisons of new models in the future.

As we move into the subsequent phases of the challenge, focusing on mortality and multimorbidity prediction, it will be crucial to consider how the insights gained from chronological age prediction can be translated to these more complex, clinically relevant outcomes. The diverse approaches showcased in this first phase provide a strong foundation for tackling these challenges, potentially leading to more accurate and personalized risk assessments for age-related diseases.

## Methods

### Challenge Design and Participation

The Biomarkers of Aging Challenge was structured in three phases: Chronological Age Prediction (March 1 - July 5, 2024), Mortality Prediction (July 1 - November 1, 2024), and Multimorbidity Prediction (scheduled for 2025). This study focuses on Phase 1 results. Teams could submit daily predictions, with the best result retained for final judging. For Phase 1, finalist submissions needed to achieve a mean absolute error (MAE) of ≤3 years between predicted and actual ages.

### Dataset Generation and Preprocessing

We generated a novel dataset using the Illumina MethylationEPIC v2.0 platform, comprising DNA methylation profiles from 500 blood samples provided by Mass General Brigham Biobank. The dataset covered ages 18-99 years and was balanced for sex. Participants received this data without phenotypic information for model development and testing. For training, teams were allowed to use any public or private datasets not included in the scoring set. The Biolearn platform provided access to relevant data from Gene Expression Omnibus (GEO), National Health and Nutrition Examination Survey (NHANES), and Framington Heart Study (FHS), though its use was optional.

### Evaluation Criteria

For Phase 1, submissions were evaluated using MAE on a held-out test set. A public leaderboard, updated in real-time using 50% of the data samples, aided teams in model development. Final rankings were determined using the full dataset.

Since DNA methylation data has limited precision, we considered any MAE within 0.1 to be tied with ties broken in favor of the model using the least number of features. Specifically, for first place, the best MAE score was considered against every other score within 0.1, and the model using the least number of features was the winning model. Second place was determined in the same way, with the first place model score removed and third place with the first and second place models removed.

### Phase 1 Winning Model Architectures

The first-ranked entry’s Skip-Improved Training Hive (SITH) Network: This ensemble model comprised multiple feed-forward (FF) neural networks, each with a linear skip layer. It utilized 8,000 CpG sites from 19 datasets (>8,000 samples total). Each base model (∼500,000 parameters) consisted of three hidden layers with 64 neurons each and ReLU activation. The linear skip layer connected the input directly to the output neuron. The ensemble’s final prediction was the unweighted mean of all base model predictions. Training employed the AdamW optimizer with OneCycle scheduling over 3000 epochs and an age-weighted smoothed-L1-loss function, which required a modest 4 GB of GPU memory on a Nvidia T100 for around 8 hours. To account for batch effects, each base model was trained on a subset of the data with one dataset removed and 90% of the remaining samples randomly chosen. Validation used three independent datasets (∼1,000 samples). The SITH network achieved an MAE of 2.45 on the full competition dataset.

The second-ranked entry’s CpGPT (CpG Pre-trained Transformer): This transformer-based model flexibly used a subset of CpG sites from a vocabulary of 5,773 probes derived from 26 existing epigenetic clocks. The model architecture included a chromosome encoder, genomic position encoder, beta value encoder, transformer encoder, two beta value decoders, an age decoder, and a covariates decoder, totaling approximately 1.6 million trainable parameters. CpGPT was pre-trained on 105,850 samples from 1,566 GEO datasets for about 1,000 epochs using 4 NVIDIA A10G GPUs over ∼14 days. It was then fine-tuned on ComputAgeBench data for 30 epochs. The model demonstrated high stability, with age predictions across different input CpG subsets showing correlations consistently above 0.999. For the competition, CpGPT achieved an MAE of 2.55 years.

The third-ranked entry’s Deep Learning ResNet: This dense ResNet architecture processed 136,111 CpG sites from 13,446 samples across 33 GEO datasets (mean age 50.2 years). Preprocessing involved mean-imputation of normalized beta values, exclusion of cross-reactive probes and sites with large variances, and adding 0.5 years to integer age labels. The model structure included an initial fully connected layer, three residual blocks (each with two fully connected layers and ReLU activation), and a final fully connected output layer. Training used a balanced split of young (20-24 years), old (>95 years), and middle-aged (24-95 years) individuals. The Adam optimizer was employed with hyperparameters optimized through random search. The training ran for 500 epochs with a batch size of 64 on an NVIDIA A100 GPU, using Mean Squared Error as the loss function. Linear regression was applied post-training to reduce potential linear bias. The model achieved an MAE of 2.46 on the competition dataset.

### Statistical Analysis

All statistical analyses were performed using standard Python libraries. The code for these analyses will be made open-source, ensuring reproducibility and transparency of our results.

## Data and Code Availability

The scoring DNA methylation dataset has been released as an open-access resource following challenge completion at GSE246337^23^, while the phenotypic data will only be released after completion of all challenge phases. The Biolearn platform, including standardized implementations of existing biomarkers and evaluation tools, is available at https://github.com/biolearn/biolearn. The workflow for the challenge is available at https://github.com/biolearn/biomarkers-of-aging-challenge-2024. All analysis code for this paper will be made available through the same repository.

## Acknowledgements

We thank all participants in the Biomarkers of Aging Challenge for their contributions. This work was supported by the Volo Foundation and Methuselah Foundation, the National Institute on Aging (NIA), the Hevolution Foundation, and Bob Rosencrantz. We acknowledge the Mass General Brigham Biobank participants and staff for the samples used in this study. KY was supported by NIA F99AG088431.

## Author Contributions

KY, SP, DB, JRP, MM, and VNG designed the challenge. JR, LPC, JT, and SJ participated in the challenge and contributed to the winning models. KY and SP analyzed the data. KY drafted the manuscript. All authors reviewed and contributed to the manuscript. MM and VNG supervised the study.

